# AVATAR: AI Vision Analysis for Three-dimensional Action in Real-time

**DOI:** 10.1101/2021.12.31.474634

**Authors:** Dae-Gun Kim, Anna Shin, Yong-Cheol Jeong, Seahyung Park, Daesoo Kim

## Abstract

Artificial intelligence (AI) is an emerging tool for high-resolution behavioural analysis and conduction of human-free behavioural experiments. Here, we applied an AI-based system, AVATAR, which automatically virtualises 3D motions from the detection of 9 body parts. This allows quantification, classification and detection of specific action sequences in real-time and facilitates closed-loop manipulation, triggered by the onset of specific behaviours, in freely moving mice.

## Main

AI-based tools for high-throughput analysis of human and animal behavioural analysis have much gained popularity recently. These have included the use of Convolutional neural networks (CNN) using markerless video recordings from a single camera [1-5] and reconstruction of 3D skeletons using multi-camera systems [6-10]. However, the creation of a system that can record, quantify, classify and detect specific behaviours in real-time has been challenging. To address this, we established AVATAR (AI Vision Analysis for Three-dimensional Action in Real-time), which can detect and reconstruct an animal’s body movements in 3D and is capable of detecting behaviours in real-time to facilitate closed-loop manipulations in freely-moving, markerless mice.

To obtain image data from multiple angles, we designed a chamber, “AVATAR studio” (Fig.1a; Supplementary Fig.1,2; and Methods); consisting of five Full-HD CMOS cameras (1200 × 1200 pixels, 30 frames per second) for recording a moving subject at different viewpoints (1 bottom + 4 side views), LED lights device for preventing motion blur in a transparent box. The bottom panel is made transparent to visualise limbs under the body. We confirmed that key body areas: the nose, head-top, tail-tip, and four paws, can be monitored by at least two of the five cameras at any given time, which enables their co-ordinates to be calculated. To prevent time consumption, due to sequential search of images generated by five multi cameras, image data from the five cameras were concatenated into a single collage (Supplementary Fig.2). An object detection algorithm [11, 12], was modified to search for major body parts on mice. The network, called “AVATARnet”, can detect 9 body parts in a collage and infers the center point of each object within 30 ms (Fig.1b). Based on the yolov4 network, AVATARnet has modified the input network size (608×608 pixels) and number of parameters (56.9M), for detecting small body parts. We also added data augmented methods that randomly change the orientation of body parts (Methods) (Fig.1b, Supplementary Fig.3).

We trained AVATARnet with labeled data (C57BL/6J, ICR/J mouse) from over 1,000 images captured by AVATAR studio under various conditions; solitary, with a conspecific, or with a non-social object. The trained AVATARnet successfully detected the 2D position (x-y) of body parts in any video data obtained from the five cameras at a level comparable to that of human labeling (center point difference = SD < 4.04 ± 3.71 mm). (Fig.1c; Supplementary Fig.4, 5; Methods; and Supplementary Video 1). We devised a posture set algorithm and corrected for detection of errors or wrong paw directions (left, right) during body object detection.

Next, we calculated the 3D coordinates of body parts from the 2D information obtained by AVATARnet. We used a triangulation and bundle-adjustment algorithm [13] (Fig.1d) to calculate the 3D position of particular body parts using the 2D video data obtained from at least two cameras, camera-intrinsic parameters (focal length, principal point, skew coefficient) and camera-extrinsic parameters (the location and orientation of the camera). We reconstructed virtual motion by connecting the 3D coordinates of nine body parts with eight vector sets, “action skeletons”, that ran along the skeletal structure of a mouse (Fig.1e; Supplementary Fig.6,7; and Methods). The virtually-reconstructed “AVATAR mice” are action skeletons representing whole-body motions in 3D virtual space (Supplementary Video 2). Finally, through the AVATAR system, 3D coordinates of each body-part over every frame (20fps) can be obtained and analysed in various ways (Supplementary Fig.8).

We tested the motion capture ability of AVATAR in various states. AVATAR successfully detects dystonia in tottering mice (a1A^tot/tot^) [14], characterised by twisted body postures, and can record freely-moving mice with a head-mounted optic wire. It can capture the individual motion of two mice housed together and shows no interference in the presence of human-made objects. (Supplementary Fig. 9; Supplementary Video 3).

AVATAR revealed that while a mouse could yield more than 2,000 unique poses (i.e., non-redundant pose sets) during 5 minutes of exploration in the chamber, the motion complexity obtained from 50 C57BL/6J mice showed a higher number during the same time (53,397 unique motion units, total frames = 450,000; confidence interval, ±5 mm) (Supplementary Fig. 10, 11 and Methods),

All the behaviours of the freely moving mice are quantified with 3D skeletal coordinates, which can be converted into a time series (Fig.1f). Skeletal coordinates can be parameterized according to various criteria and the time period can be classified according to specific rules. In this paper, we classified the AVATAR mice behaviour over time using an ethogram. Since the value of the skeletal coordinates varies across episodes of behaviour, depending on the orientation of the mouse, the spine skeleton of the mouse was fixed to the x-axis to align the orientation across all episodes (Fig.1g). Through human labeling, the exploratory behaviour of mice was classified into five major ethograms (walking, sniffing, rearing, immobility, grooming) in AVATAR studio (training set: 18000 frame) (Fig.1h). We designed an LSTM neural network classifier [15] to classify behavioural time series data derived from the AVATAR system (Fig.1i). The LSTM classifier training accuracy and training loss converge to 89.91% and 0.27, respectively at 100 epochs (Fig.1j). LSTM annotation prediction performed with an error rate of 11.97% when compared to human-labelled data (Fig.1k, l, m). Additionally, the training dataset can undergo further unsupervised clustering to produce finer classifications (Fig.1n).

To test the efficiency of AVATAR in real-time analysis and feedback, we set up a system for closed-loop photostimulation in response to TTL signals generated by the AVATAR system during specific motion sequences. To this end, we unilaterally injected 0.5 µl of AAV2/9-Ef1α-dflox-hChR2 (H134R)- mCherry or control gene constructs (Addgene, USA) in the VTA (AP, -3.1 mm; ML, 0.4 mm; DV, – 4.5 mm, from bregma) in dopamine transporter::Cre (DAT-Cre) mice with a fiber-optic cannula (200 µm diameter; Doric Lenses, Canada) implanted over the VTA (Fig. 2a).

AVATAR was capable of driving photostimulation within 90 milliseconds from when mice displayed the selected motion sequence (Fig. 2b). In this paper, the rearing pose sequence (10 frames) was selected as a trigger for photostimulation (Fig. 2c, d). During photostimulation, mice showed a different pose sequence during rearing. A total of 109 rearing episodes were observed in 6 trials of 3 mice (Off: 40, On: 69), and this was composed of a time-bin of 100 frames centered on when the nose was at maximum height. Closed-loop photostimulation increased rearing time (Off: mean ± SD = 2.37±0.614, On: mean ± SD = 2.25±1.42 ms, p value < 0.05), decreased average rearing height (Off: mean ± SD = 4.19±0.946, On: mean ± SD = 2.89±0.78, p value < 0.05) and decreased peak nose height (Fig. 2d, e, f). Stereotyped up-and-down movement along the y-x plane was observed in control conditions, but showed a more complex trajectory during photostimulation (Fig. 2g, h) (Supplementary Video 4).

The AVATAR system provides tools to simulate quantitative behavioural data. The web-based simulator allows analysis and visualisation of data with various functions in the simulator (access: http://demoavataranalysis.ap.ngrok.io/) (Supplementary Video 5). Body orientation during all rearing episodes were aligned and fixed on the x-axis, which allows a stacked display of all poses during rearing episodes (Fig. 2.i).

With a set of stacked poses, changes in the mice’s nose movement were recorded on the x-z plane, and histograms of nose coordinates were drawn for each axis (Fig. 2.j). The results show alterations in the coordinate distribution of the nose during photostimulation. By observing individual rearing trials in the simulator (Supplementary Video 6), we confirmed that mice typically stopped, before raising their head and immediately descending, whereas during photostimulation, mice lift their head concurrently while moving (Fig. 2.k). Without fixing the pose to the x-axis, we visualised the entire pose sequence history in the AVATAR chamber. Rearing was mainly observed in the corners of the chambers, whereas during photostimulation, rearing was more scattered across the entire chamber (Fig. 2.l).

We compared the rearing sequence between control and photostimulated conditions by machine learning tools. Clustering with t-sne [16] showed that rearing in photostimulated mice has different features from normal rearing (Fig. 2m). A Gaussian SVM classification learner was able to distinguish between non-photostimulated and photostimulated rearing with a probability of over 97% (Fig. 2n). These results suggest that closed loop stimulation with AVATAR induces an altered rearing sequence.

Machine learning has emerged as an essential tool for the precise and unbiased evaluation of animal behaviour [17-22]. By mimicking the brain-like motion detection postulated in the biological motion theory and utilizing available video processing technologies, the AVATAR system performs real-time motion tracking of nine body parts from freely-moving mice with millisecond resolution (Fig. 1a, b) by translating a large amount of multi-viewpoint video data and reconstructing it into an “AVATAR mouse”. The virtual image consists of co-ordinates for each body part (x, y, z) while using a significantly lower amount of data (0.35 megabytes vs. 1,000 megabytes per min or 500 megabytes vs. 1440 gigabytes per 24 hours/mouse) (Fig. 1c-e). This facilitates 3D motion sequencing, clustering and action phylogeny (Supplementary Fig 13) and can be used for real-time feedback control of brain circuits through closed-loop optogenetics during specific behavioural sequences (Fig. 2a-c). Typically, closed-loop manipulations have relied on simple analog signals such as lever pressing or licking, whereas the ability of AVATAR to detect specific behaviours in real-time increases the scope and flexibility of behaviour-induced manipulations. Combining AVATAR with conventional imaging experiments allows researchers to identify behavioural correlates of neural activity, with a high degree of spatiotemporal resolution in freely-moving mice. The ability of AVATAR to detect dystonia in tottering mice (Supplementary Video. 3) demonstrates its potential in high-throughput screening of motor disorders. This can be further expanded to large-scale pre-clinical screening of drugs for such neurological disorders, as assessed by changes in 3D body motion analysis. Thus, we expect AVATAR to be a versatile tool in high-throughput analysis and manipulation of behaviour in both research and preclinical settings.

**Figure 1.**
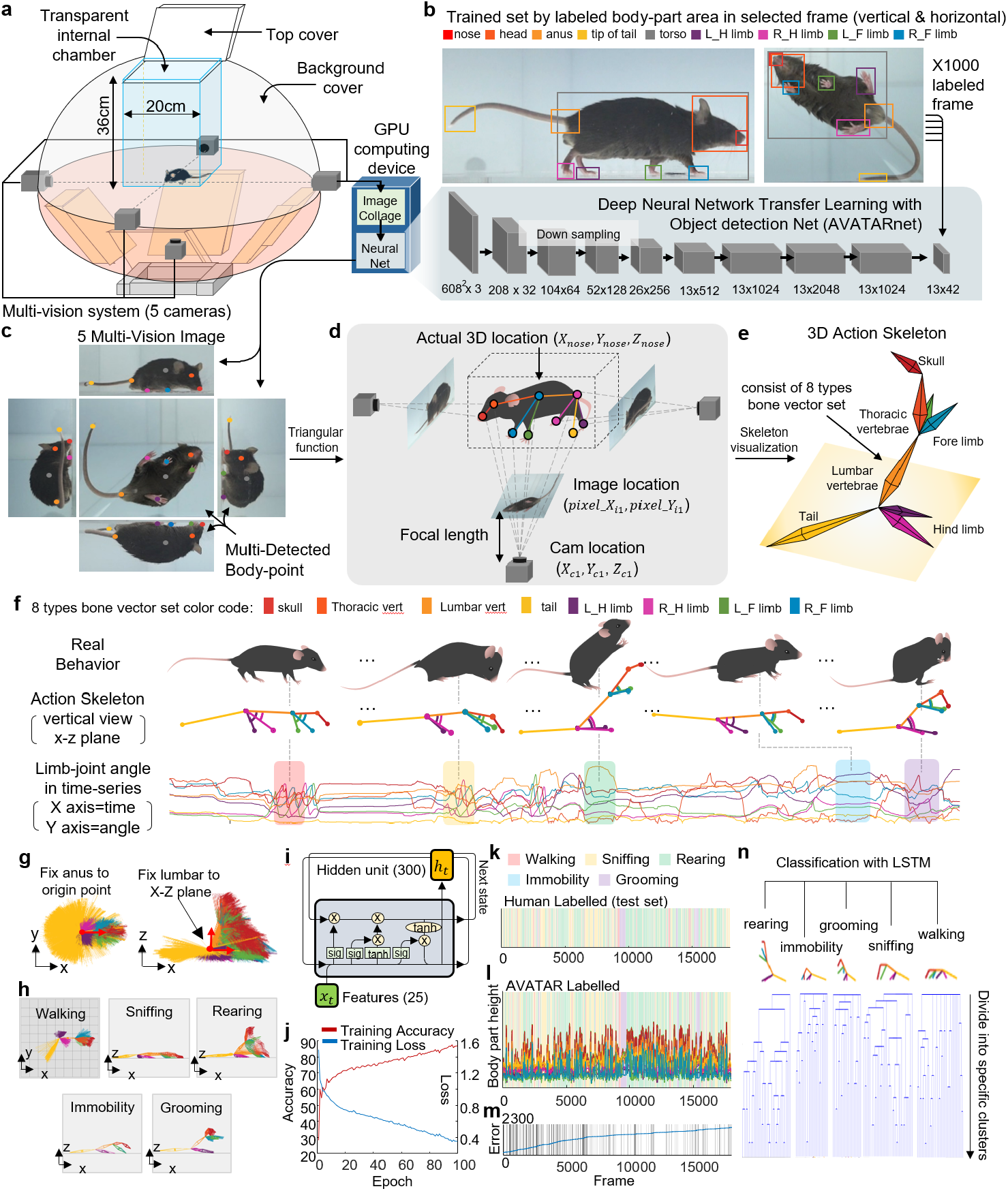
3D behavior analysis methods of AVATAR system. a. Schematic overview of the AVATAR studio, with a five-camera recording system. The mice are placed in a transparent internal chamber in the center of the studio. b. Top, each colored square box shows selected human-labeled body part areas; data sets for these labeled body parts were used to train the object-detection neural network (AVATARnet). Bottom, the network structure of the AVATARnet. c. Each body part was detected (center position) from collaged images captured by multiple cameras. d. Conceptual scheme for 3D reconstruction from multiple images in AVATAR studio. Here, only three cameras are presented for convenience. e. Reconstructed 3D action skeleton consisting of eight vector sets. Each color shows a body vector connected to the matching body parts. f. Representative behaviour sequences. Top, schematic figure of behaviour from a real mouse. Middle, action skeletons from a vertical view according to each mouse’s behaviour. Bottom, limb combination (joint angle variation) in time-series. Dashed lines indicate the behaviours and the action skeletons at the matched points. g. Left, horizontal view of stacked poses set by fixing the lumbar to the x-z plane from the origin (anus: 0, 0, 0; torso center: x, 0, z). Right, vertical view of the stacked poses. h. Visualisation of the stacked pose sequences for a specific time-bin (red arrow, walking; yellow arrow, sniffing; green arrow, rearing; blue arrow, immobility; purple arrow, grooming). i. Schematic of the AVATAR LSTM classifier model. j. Training accuracy and loss of AVATAR LSTM. k. Comparison of human annotation of exploration behaviour in AVATAR studio l. and AVATAR labelling (red, walking; yellow, sniffing; green, rearing; blue, immobility; purple, grooming). m. Cumulative graph of the number of error frames over entire frames n. A phylogenic tree classifies poses into five predefined behaviours through semi-supervised learning. The data set may be divided into more groups according to the classification resolution.

**Figure 2.**
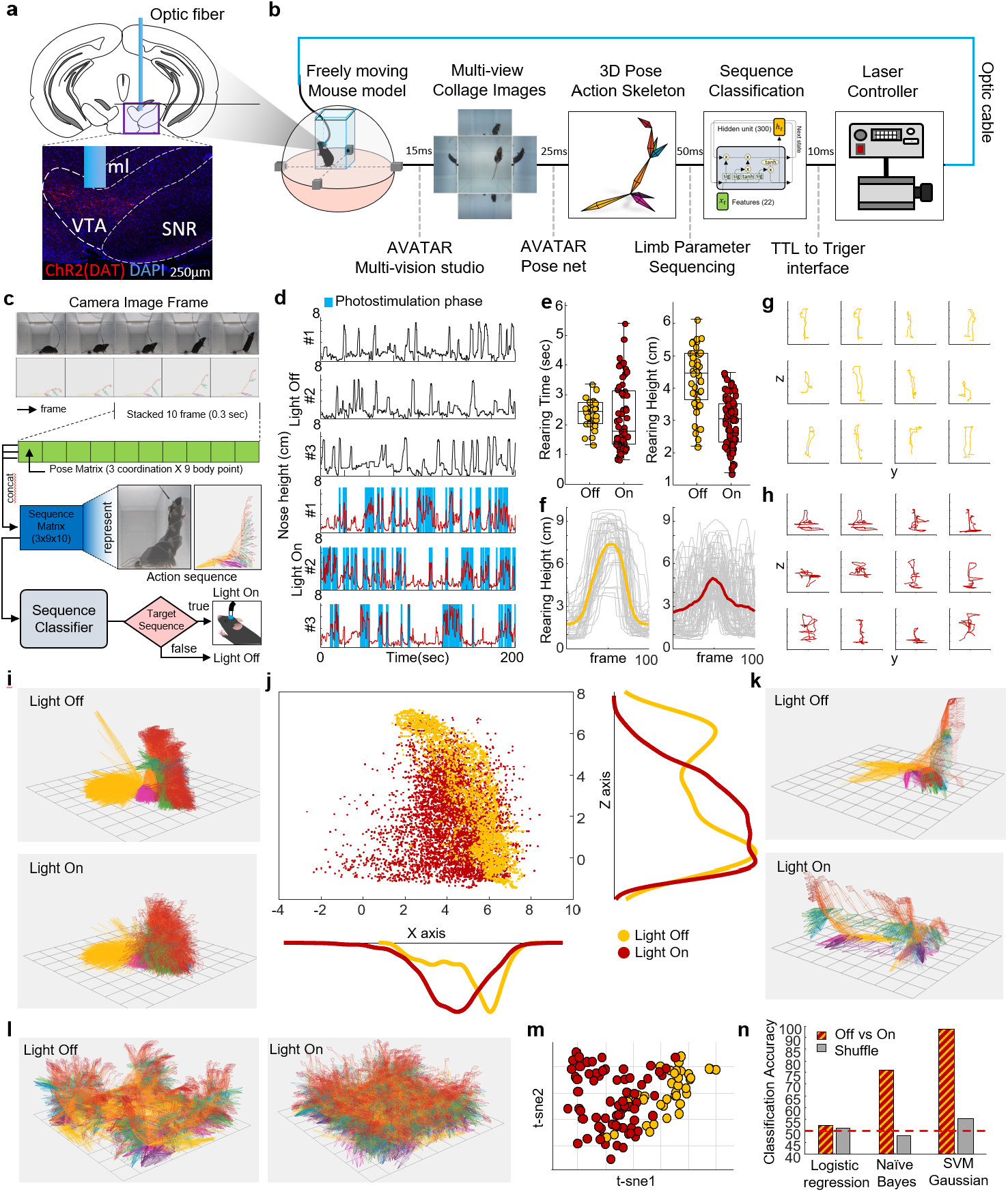
Real-time closed loop optogenetic test in AVATAR system. a. Activation of dopaminergic neurons in the VTA, by injecting 0.5 µl of AAV2/9-ef1α-dflox-hChR2 (H134R)- mCherry in the VTA of DAT-Cre mice. b. Schematic figure of AVATAR real-time closed-loop brain photostimulation. A total detection and stimulation time of 100ms is required. c. Schematic of reinforcing the rearing process. When the mouse rearing pose sequence is detected, photostimulation is triggered to enhance the rearing behaviour. d. Nose height graph for each condition (n=3). Blue boxes represent times when rearing sequences were detected. e. Left, Boxplot of rearing time. Right, Box plot of average nose height during rearing (yellow dot = photostimulation off, red dot = on). f. Change in nose height in the extracted rearing sequence (time bin = 100 frames, 2.5 seconds) g. The trajectory of the nose in the extracted rearing sequence during no photostimulation (y-z plane) h. The trajectory of the nose in the extracted rearing sequence during photostimulation (y-z plane) i. Visualisation of extracted rearing sequence poses when stacked and aligned on the x-axis. j. Histogram of the nose position projected onto the x-z plane during rearing (yellow dot = photostimulation off, red dot = on). k. Example of stacked poses during rearing (not fixed to the x axis) l. History of all poses within the AVATAR chamber m. t-sne clustering of rearing with no photostimulation and with photostimultaion (yellow dot = photostimulation off, red dot = on). g. Machine learning classifier of rearing data. Gaussian SVM predicts with 97% accuracy.

## Acknowledgements

This study was supported by a grant from Samsung Science & Technology.

## Author Contributions

D. K. designed the study and coordinated the experiments. D-G. K., Y-C. J., and A. S. performed the behavioural experiments with the AVATAR system. D-G. K. developed the AVATAR hardware and software algorithm platforms. All authors participated in writing the manuscript.

## Methods

### Animals

For wild-type experiments, 50 male C57BL/6J (Jackson Laboratories stock #000664) adult mice (8-16 weeks of age) were used. For the dystonia mouse model, three male B6. D2 ™ Cacna1α^tg^/J (Jackson Laboratories stock #000544) adult mice (8-16 weeks of age) were used. All mice were group housed at five to six mice per cage under a 12 h light/dark cycle and *ad libitum* access to food and water. All procedures were conducted according to the Korea Advanced Institute of Science and Technology (KAIST) Guidelines for the Care and Use of Laboratory Animals and were approved by the Institutional Animal Care and Use Committee (Protocol No. KA2014-05).

### Recording behaviour in the AVATAR studio

A transparent chamber was made of 5-mm-thick acrylic panels (200 mm x 200 mm x 300 mm) and inserted in the center of AVATAR studio (Supplementary Fig.1a, 2c). For video recording, five high-speed cameras (FLIR ® Systems, Inc., BFS-U3-23S3C-C) were installed at five different viewpoints: front, rear, left, right, and bottom (Supplementary Fig.1a,2a,2b). The specific positions of the cameras were adjusted according to the focal length of the camera (30 cm). The operation of the five cameras was synchronized to a central clock system and controlled by a host card (IOI Technology, U3×4- PCIE4XE101, 4port usb3.0 to PCIe x4 Gen 2 host card) (Supplementary Fig.2d). An opaque panel was used to cover the top to block external visual stimuli. The size, design and components of the AVATAR studio are detailed further in Supplementary Figure 1.

For behavioural recordings of wild-type mice (Fig.1f-h), we recorded one wild-type mouse at a time in the AVATAR studio. For behavioural recordings of a disease mouse model, we recorded one dystonia model mouse at a time in the AVATAR studio. For behavioural recording of the separated social model, we recorded two wild-type mice at a time in the AVATAR studio. All animals designated for behavioural recordings were habituated in the AVATAR studio for 10 minutes. Thereafter, all five cameras recorded for 5 min (Supplementary Fig. 2e).

Images were captured from each camera and transferred to the host card via a USB3.1 data cable. The host card was connected to the main computer through the PCI-Express slot and the image data were integrated into a multi-view collage image at the main computing processor unit (Intel, i9 9900k). It took about 30 milliseconds to receive each image data from the five cameras and combine them into one collaged image (30 fps).

### AVATARnet

To detect body parts from the multi-viewpoint image data obtained using the AVATAR studio, we developed a CNN algorithm called “AVATARnet” by modification of a previously reported deep-learning algorithm, “yolov4”, which exhibited good performance in accuracy and speed [12]. AVATARnet consists of 53 convolution neural network layers that have a higher input network size (608 × 608 fixels) compared to the previously reported algorithm and we added skip-connection and batch normalization (Supplementary Fig. 3a). These changes increased the detection performance of multiple small-sized objects (forelimbs and hindlimbs). We also added several training methods to enable accurate detection of body parts of various sizes. For general regularisation, we added a dropout method and data augmentation in the labeled training set.

### Training AVATARnet with mouse image data sets

We prepared multi-viewpoint images of 3,200 × 2,000 pixels for nine body parts (nose, head, torso, anus, tail-tip, forelimbs and hindlimbs) (Supplementary Fig. 3b) of C57BL/6J mice, and trained AVATARnet using the transfer learning method. To enhance detection ability, we prepared additional image data taken under different conditions, including hunting (200), social interaction (200), non-social object interaction (200), with a head-mounted optic fiber (100), and with a head-mounted endoscope (100). After training was performed using the basic data set (1,000 images for each body part class) obtained from solitary exploration in AVATAR chamber, we used sum-squared error between the predictions and the ground truth to calculate loss. The loss function was composed of the classification loss 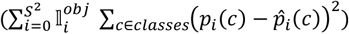, localization loss 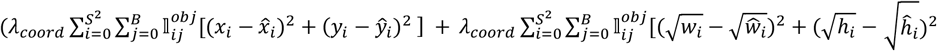, and confidence loss 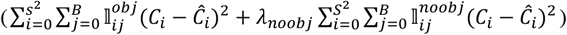. The average loss value was below 0.5 when we used more than 8000 iterations (Supplementary Fig. 3c). The training took around 12 hours; 14,000 iterations were applied using a GPU (NVIDIA, RTX 2080ti) computer processing the 3,200 × 2,000 pixel training image data set.

### Evaluation of AVATARnet with the mouse image data set

We used mean Average Precision (mAP), Intersection over Union (IOU) and Mean Square Error (MSE) to evaluate our network. IOU analysis evaluates the portion of overlay between the ground truth and a prediction box (Supplementary Fig. 4a) and our network showed more than 75% IOU after the training (Supplementary Fig. 4b). We used mAP measure to measure the accuracy of the network predictions; it evaluates the degree of overlap between the predicted body part areas and actual body part areas (Supplementary Fig. 4c) and found that AVATARnet detected the nine body parts with 90% accuracy when using mAP, which was previously used as a default metric for precision in the PascalVOC object detection competition. We also evaluated the mean square error (MSE) distance for each class (Supplementary Fig. 4d). After the network was trained, it showed a 7∼15 pixel (1.4 ∼ 4.5 mm) MSE distance from each class (Supplementary Fig. 4e, f). In addition, we confirmed that the center points of body objects detected by AVATARnet were quite similar to those marked by human observers (Supplementary Fig. 5).

### Automatic calculation of the 3D positions of body parts

To calculate the 3D position (x-y-z) of body parts from the 2D (x-y) coordinate data obtained from the five cameras (Fig.1c-e), we used a computer-vision 3D calibration and reconstruction algorithm [13]. The 3D coordinates of the target points are computed by calculating the intersection of the straight lines that pass from the center of the camera through each target point on the image (taken from focal length). We applied this algorithm to compute the 3D coordinates of the selected endpoint locations. Using the parameters of each camera (intrinsic, extrinsic and lens distortion), this method could compute the 3D coordinate of body parts from multiple 2D images (Supplementary Fig. 6).

### Reconstruction of an AVATAR mouse using an action skeleton

To make a virtual subject, the 3D coordinates of all body parts (one point for each body part) were connected with eight lines to form an “action skeleton”, which represents the vectorial location of specific skeleton parts relative to their attachments to the body trunk in 3D space (vector-1, from head to nose; vector-2, from torso to head; vector-3, from torso to anus; vector-4, from anus to tail-tip; vector- 5, from torso to left forelimb; vector-6, from torso to right forelimb; vector-7, from anus to left hind limb; vector-8, from anus to right hind limb). Unlike a real skeleton, the action skeleton is flexible in length as it represents the distance between the two connected objects. Action skeletons were generated at 20 frames per second.

### Motion unit analysis

A motion unit was defined as the set of action skeleton structures at a given time. To isolate non-redundant motion units, we eliminated redundant sets based on the confidence interval, *x* ± 1.96 × *SE* (2.5 mm) (Supplementary Fig. 8.a,b). We displayed all non-redundant motion units according to their level of frequency in a posture ring that ran counterclockwise from the 9 o’clock position (Supplementary Fig. 8.a,b). Along the time series, connecting one motion unit at the first posture ring to the next motion unit at the next posture ring yielded a behaviour cylinder that shows the dynamic changes of the motion sequences (Supplementary Fig. 8.c,d). Analysis of the behaviour cylinder revealed that there are dominant motion sequences that can be annotated as specific actions, such as walking, running and rearing. These results suggest that the AVATAR system can be useful for unsupervised characterization and quantification of motion and action sequences.

### Web simulator of the AVATAR system

We developed a web simulator for the AVATAR system. The raw file of the 3D behaviour information observed in the AVATAR studio is saved as a csv. By loading this csv file into the web simulator, the user can simulate the behaviour (Supplementary Fig. 12.a,b; Supplementary Video 4). The simulator visualises the action skeleton from a csv file containing body-part coordination; the user can observe and visualize motions frame by frame or pose sequences by setting a time-bin. In addition, each body-part can be observed separately and the history of the entire observed behaviour can be stacked.

