## Supplementary Figures for "AVATAR: AI Vision Analysis for Three-dimensional Action in Real-time"

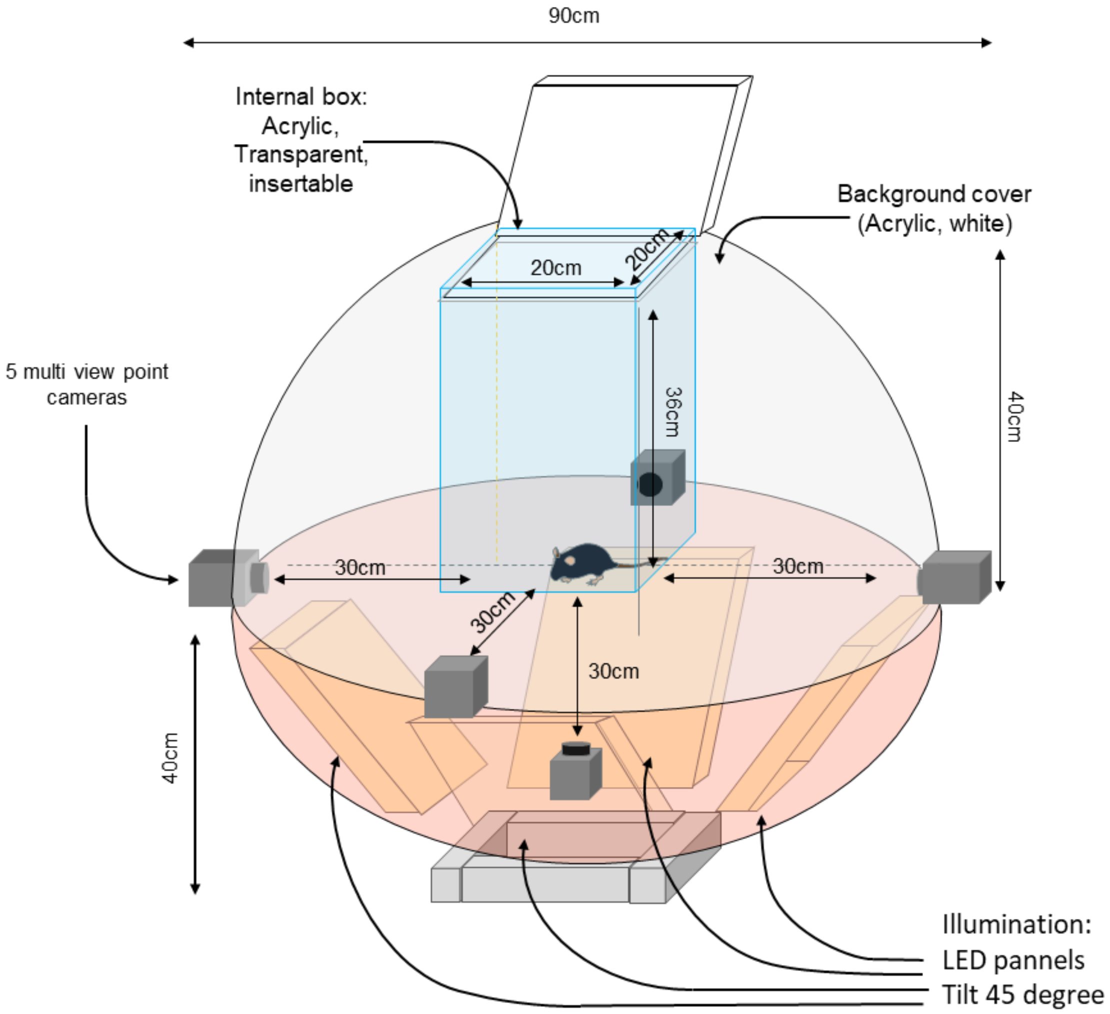


**Supplementary Figure 1: Schematic of the AVATAR studio.**
A transparent chamber made of 5-mm-thick acryl-amide panels (size: 200 mm x 200 mm x 360 mm) is mounted in the center, and five high-speed cameras are installed at different viewpoints (4 sides, 1 bottom). An LED lighting system is installed at the bottom to provide light for behavioral observation.

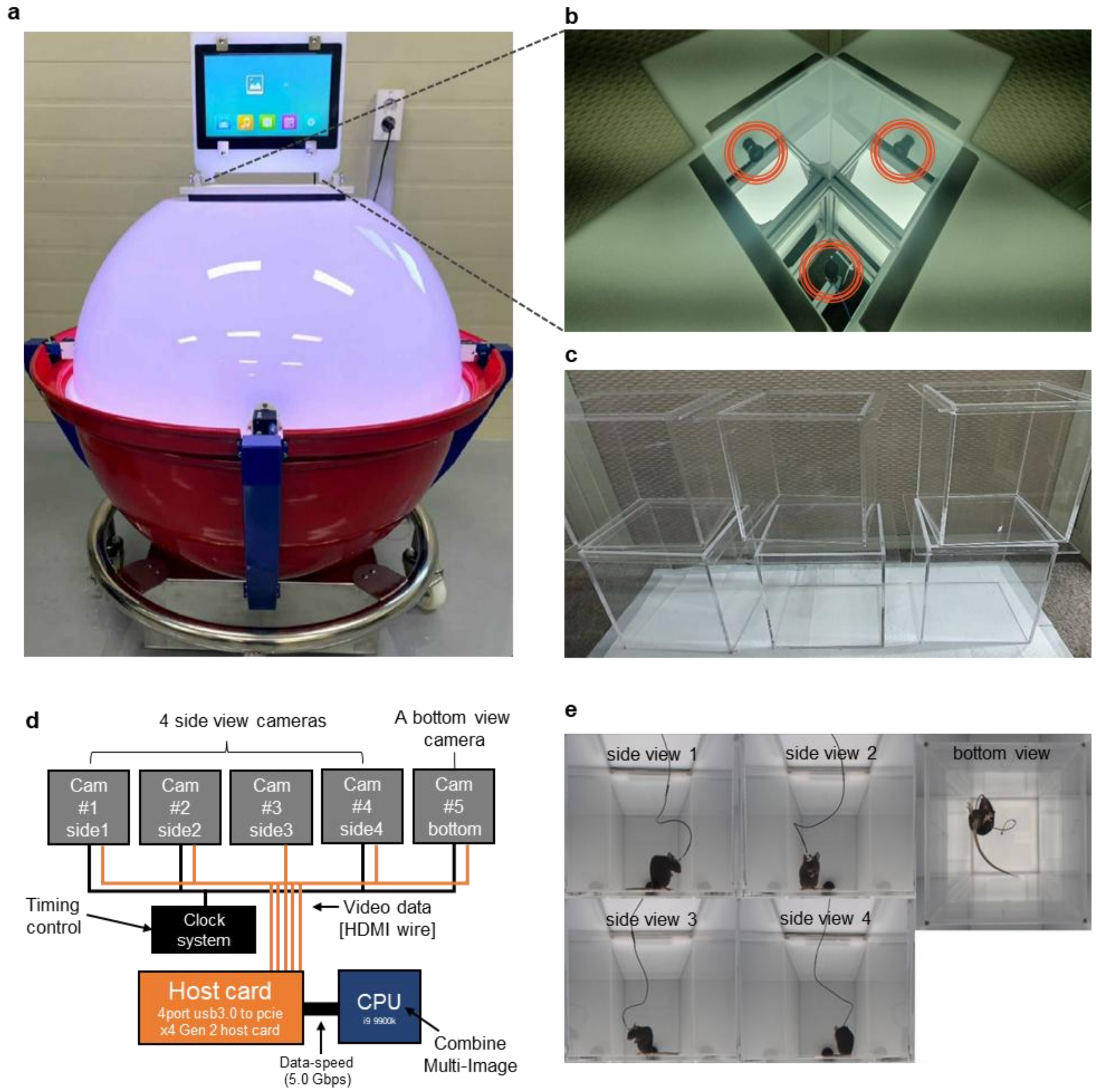


**Supplementary Figure 2: Overview of the AVATAR studio components.**
(a) Photograph of the AVATAR studio. black dot-line shows the entrance to the transparent chamber. (b) Top view of the transparent chamber. Red circles show the positions of cameras. (c) Multiple internal transparent chambers can be operated to assess multiple mice. (d) Schematic workflow of the camera network system. The five cameras are connected to the PCIe board of the PC through a USB3 cable to transfer image data. (e) An example of the multiple images captured from the AVATAR studio cameras.

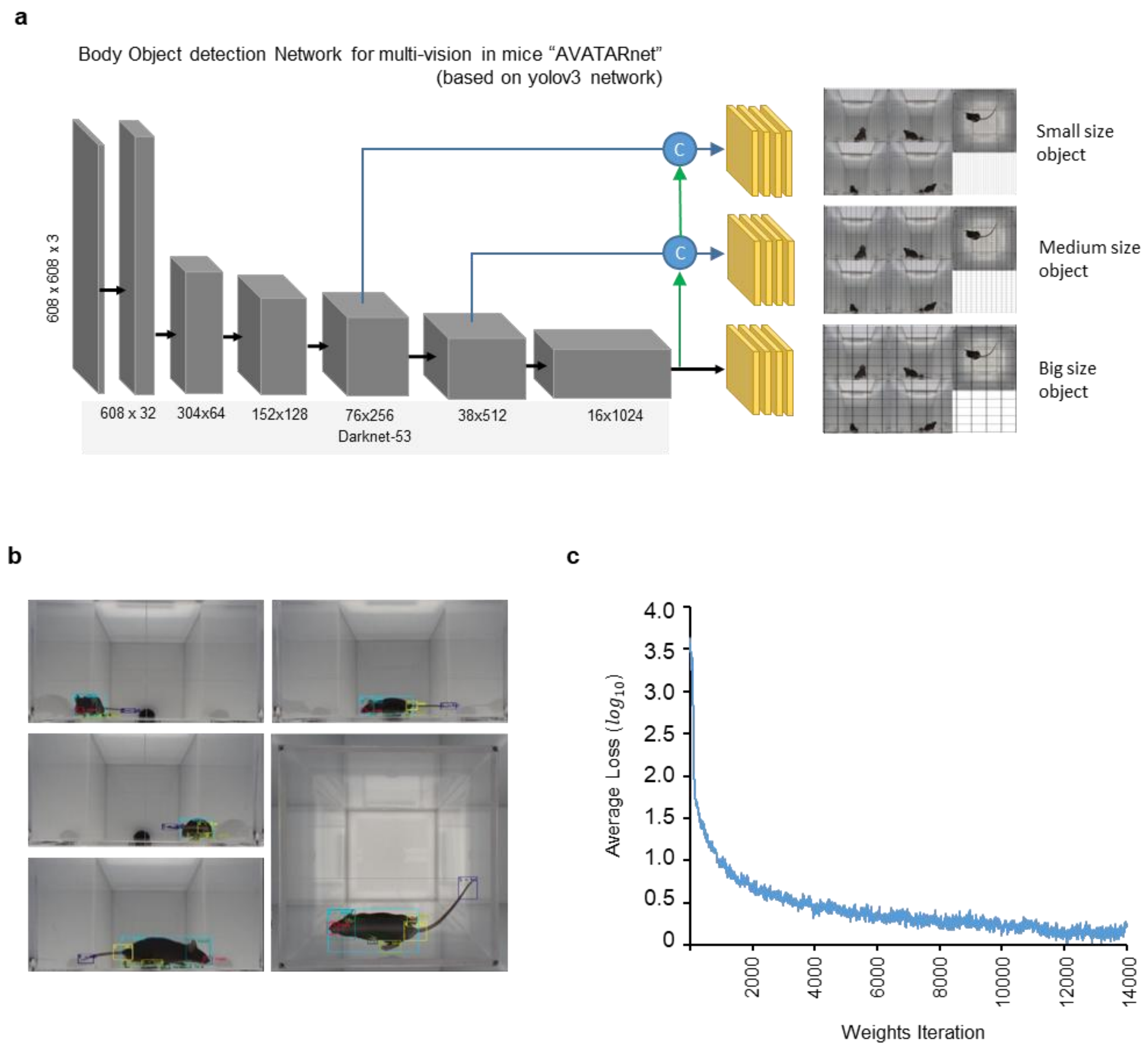


**Supplementary Figure 3: Structure and training process in AVATAR network (AVATARnet).**
(a) Structure of the AVATARnet. (b) Example of training data labeling set. (c) Training accuracy of the AVATARnet.

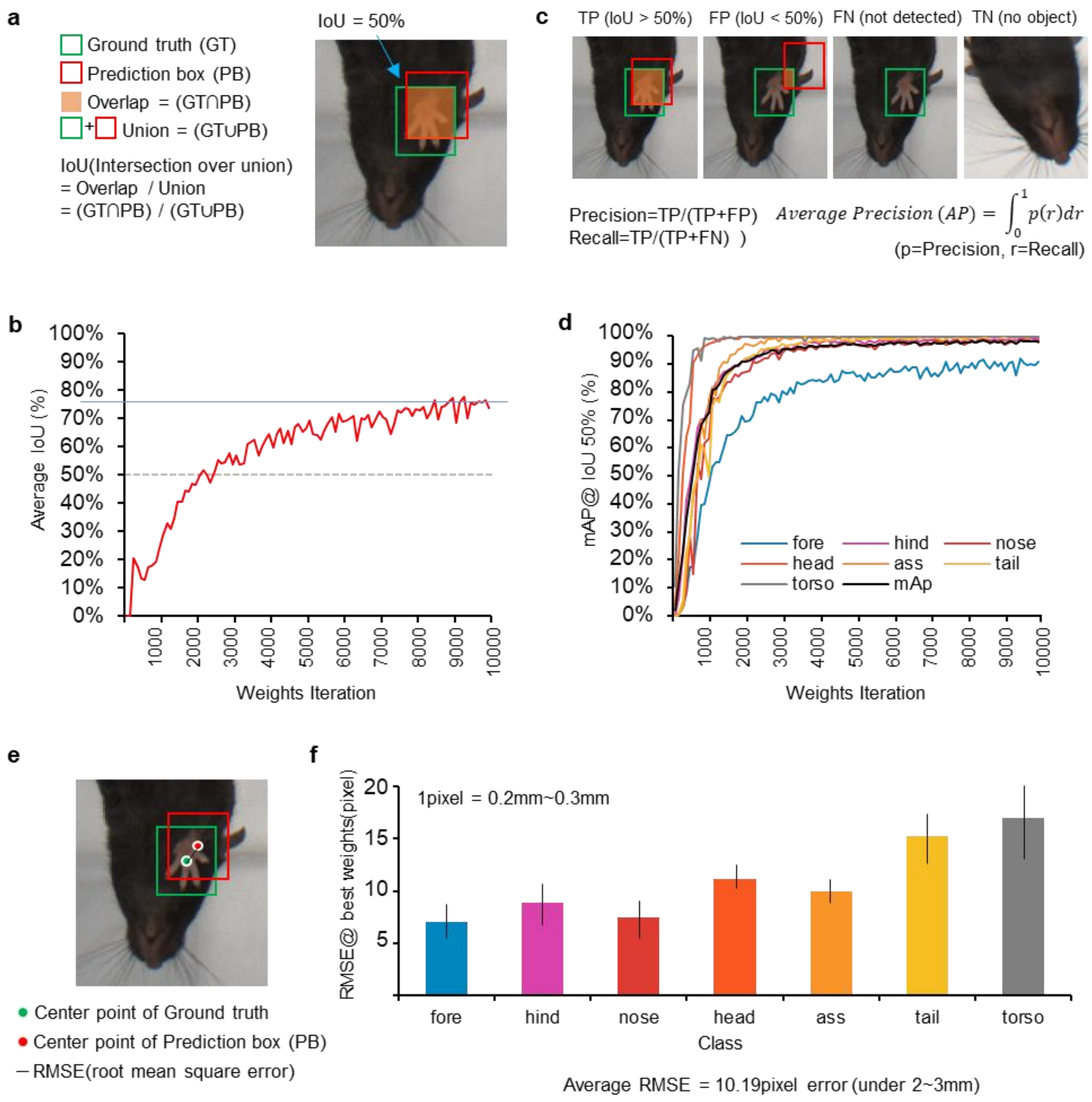


**Supplementary Figure 4: Performance evaluation with human labeling.**
(a) Description of the IOU (intersection over union) metric. (b) Average IOU progress over weights iteration. (c) Description of precision and recall. (d) Mean Average Precision (mAP) progress at 50% IOU over weights iteration. (e) Description of the RMSE. (f) Average RMSE for each body-part class.

a

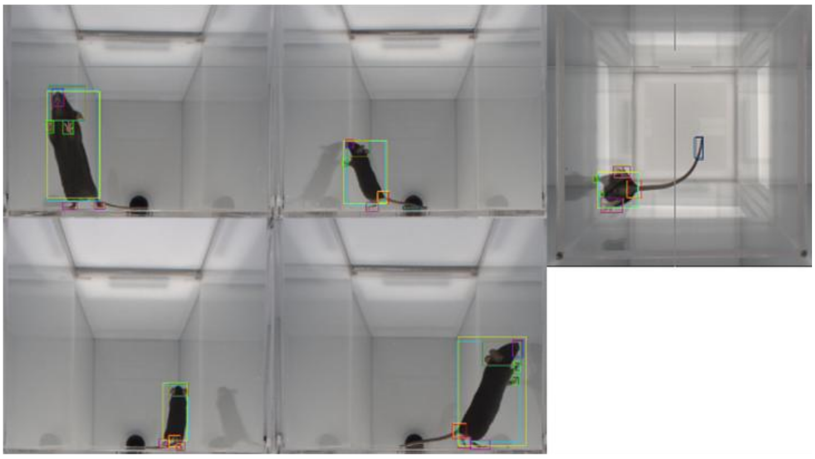

b

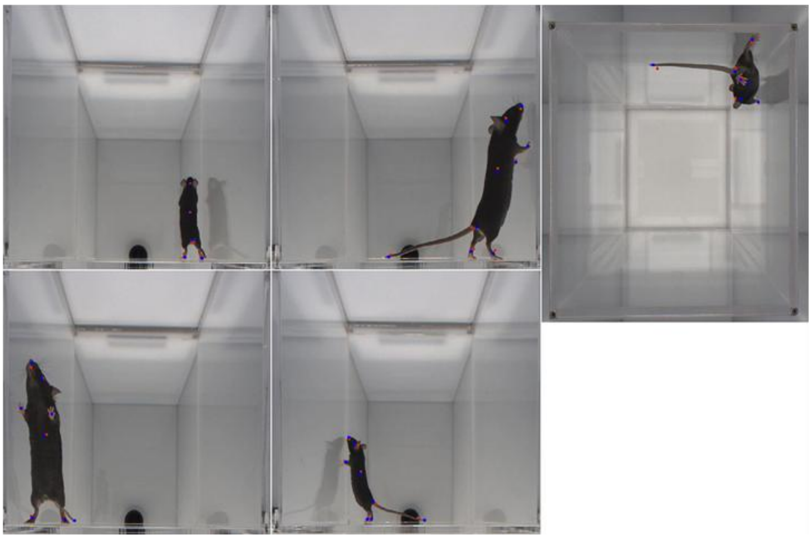

**Supplementary Figure 5: Example for comparison with human labeling.**
(a) Comparison of the body-part areas inferred by AVATARnet versus human analysis. (b) Comparison of the center points of the body parts inferred by AVATARnet versus human analysis (red dots for AVATAR; blue dots for human).

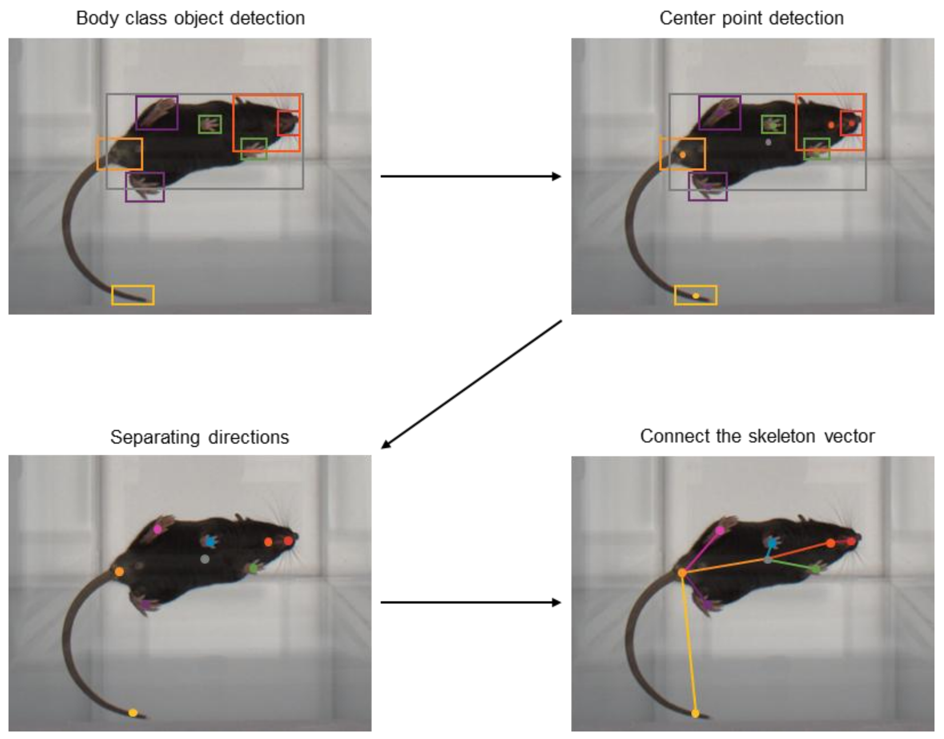


**Supplementary Figure 6: Schematic design for pose selection algorithm.**
To make a virtual subject, the 3D coordinates of all body parts (one point for each body part) were connected with eight lines to form an “action skeleton”, which represents the vectorial locations of specific skeleton parts relative to their attachments to the body trunk in 3D space. Unlike a real skeleton, the action skeleton is flexible in length as it represents the distance between the two connected objects.

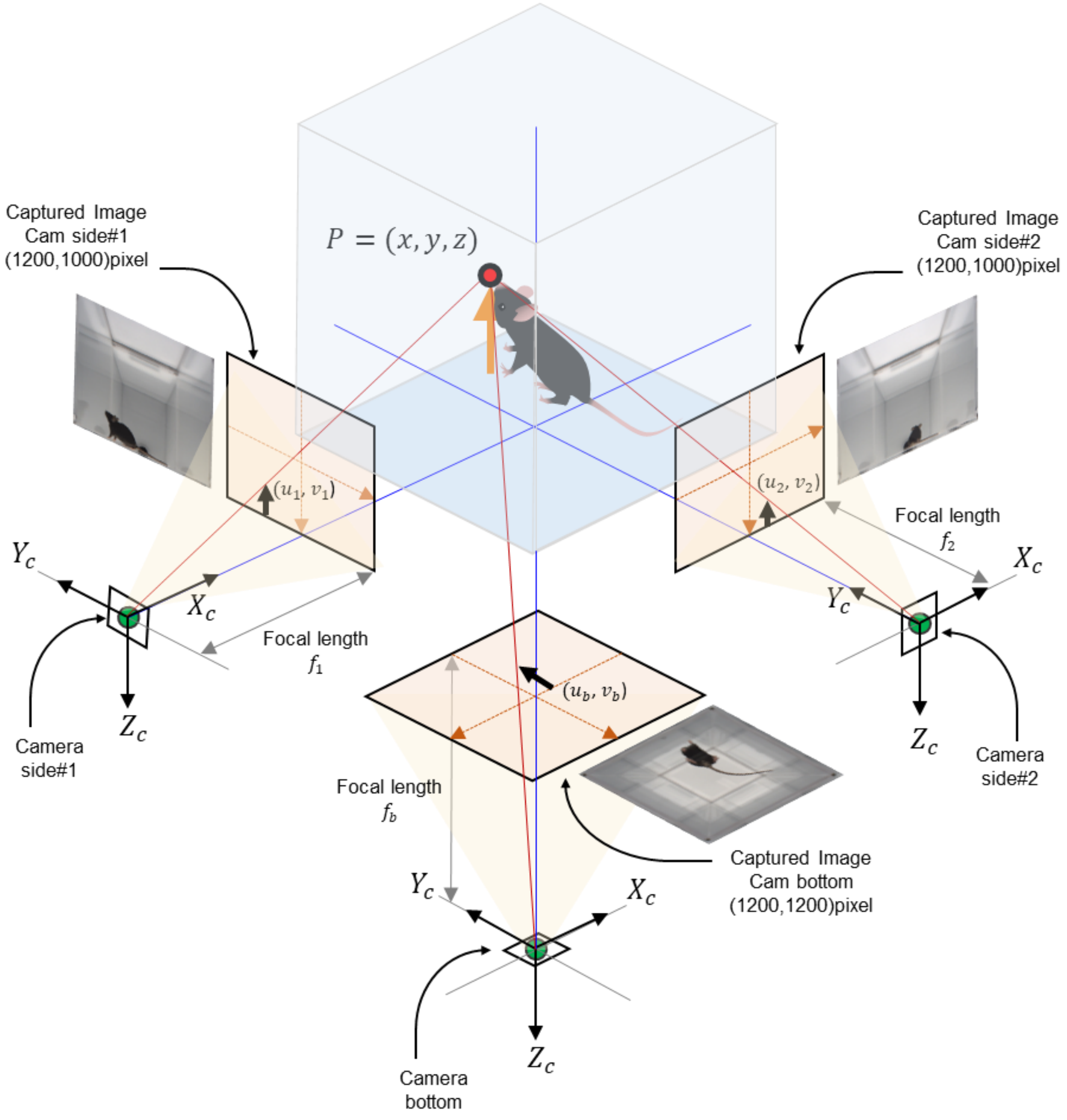


**Supplementary Figure 7: Schematic design of 3D reconstruction.**
Virtual space and positional information of the AVATAR studio. Using camera parameters of the five cameras (intrinsic, extrinsic, and lens distortion), the system can compute the 3D action skeleton from the 2D image information.

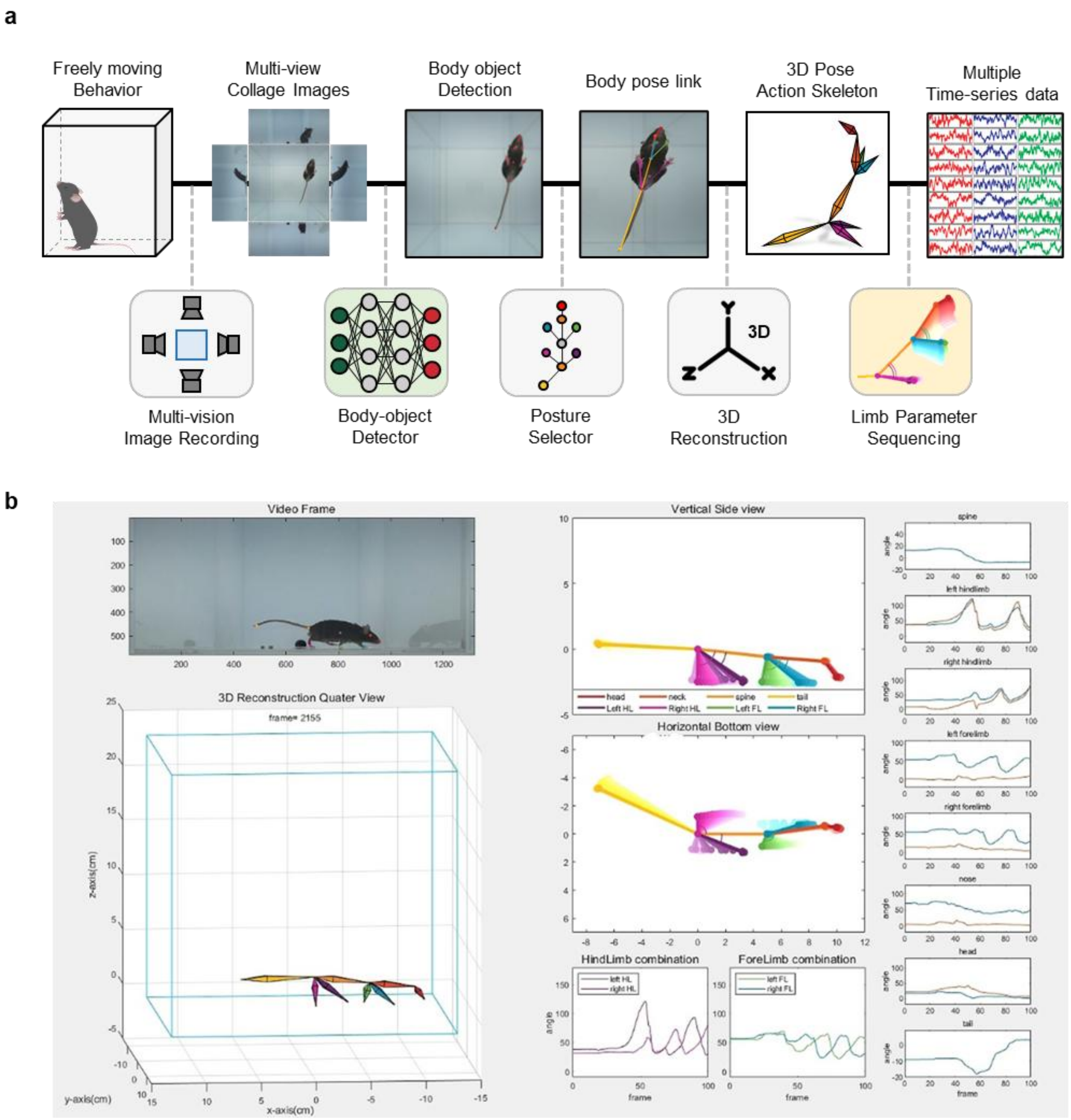


**Supplementary Figure 8: Flow chart of the AVATAR 3D analysis system.**
(a) The entire work process of the AVATAR system. (b) A representative GUI image showing real-time visualization of 3D skeletons and time-series analysis of detailed parameters for a mouse placed in the AVATAR studio.

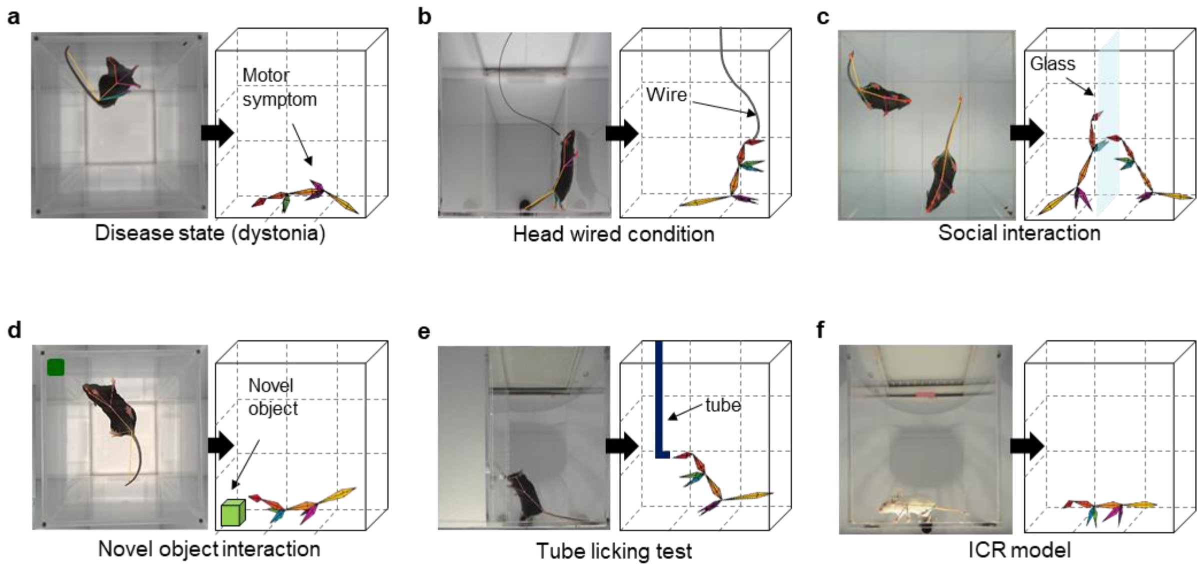


**Supplementary Figure 9: Application of the AVATAR system in various open filed behavioral experiment paradigms.**
(a-f) AVATAR analysis applied in disease model (a), head wired condition (b; for optogenetics, photometry, etc.), social interaction test (c), novel object test (d), tube test (e), and ICR mouse model (f).

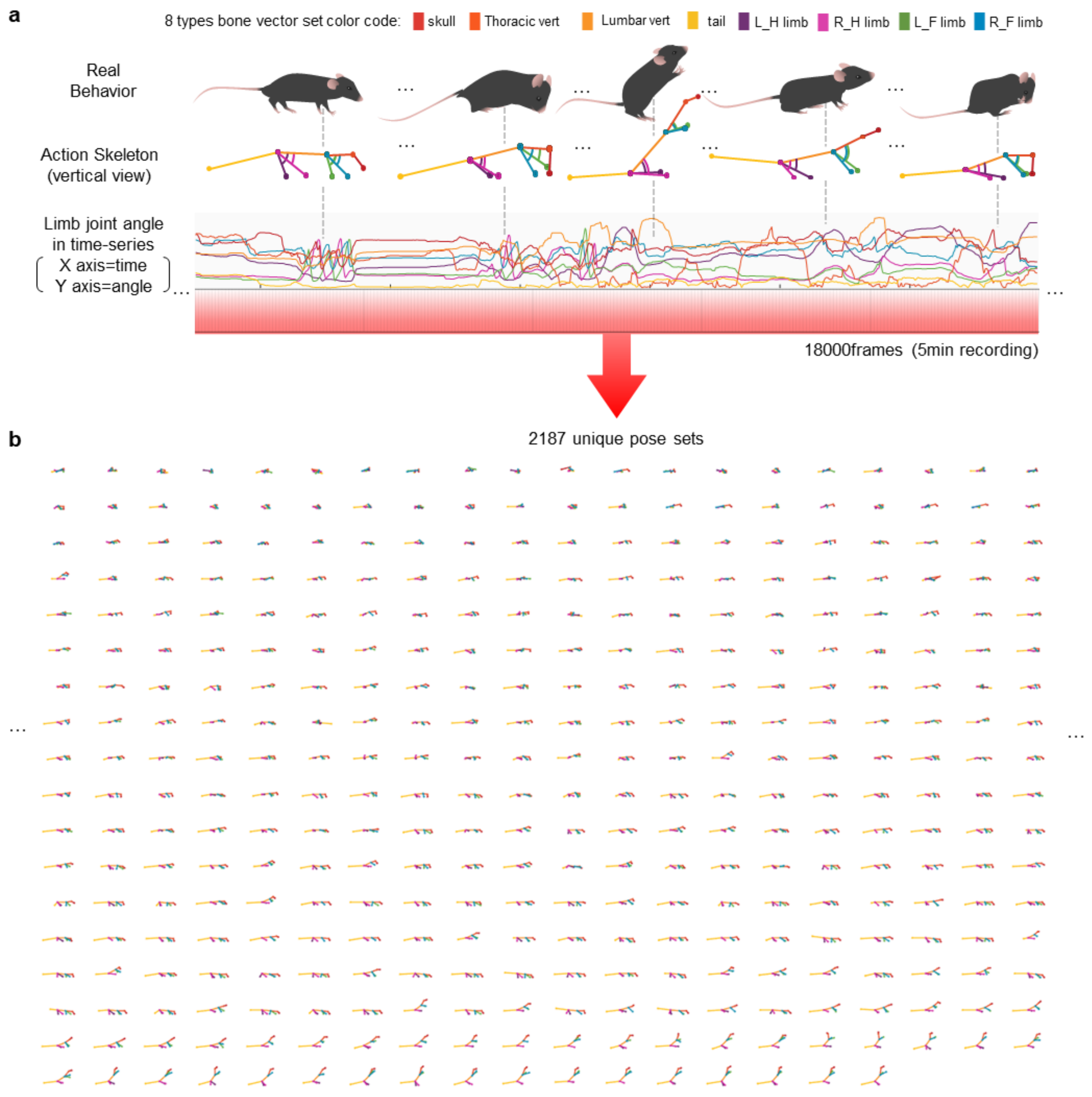


**Supplementary Figure 10: Representative classified motion units.**
(a) Time-series graph of limb joint information obtained from mice observed for 5 minutes.
(b) Eighteen thousand frames of behavioral information observed for 5 minutes can be compressed into 2,187 different unique pose sets called “motion units”.

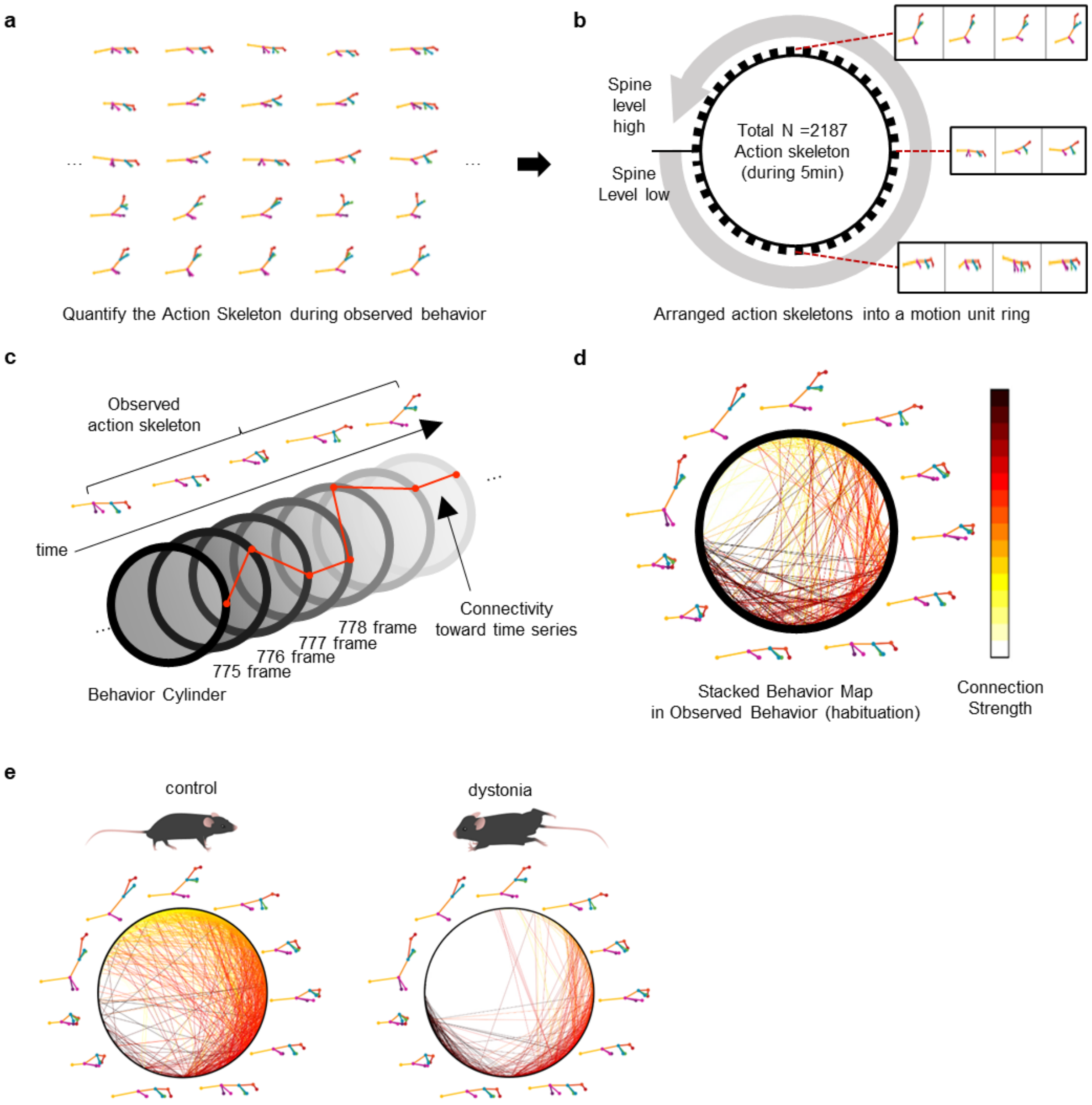


**Supplementary Figure 11: Schematic figure of stacked behavior map (the variation of pose transition).**

(a) The overall motion units reconstructed by AVATAR system. (b) Motion unit ring representing a series of motion units aligned based on similarity. (c) A motion cylinder made by stacking motion rings along with time. (d) Stacked behavior map showing the motion units linked closely (e) Comparison of wild-type and dystonia model mouse behavior in a stacked behavior map made from 5-min video data.

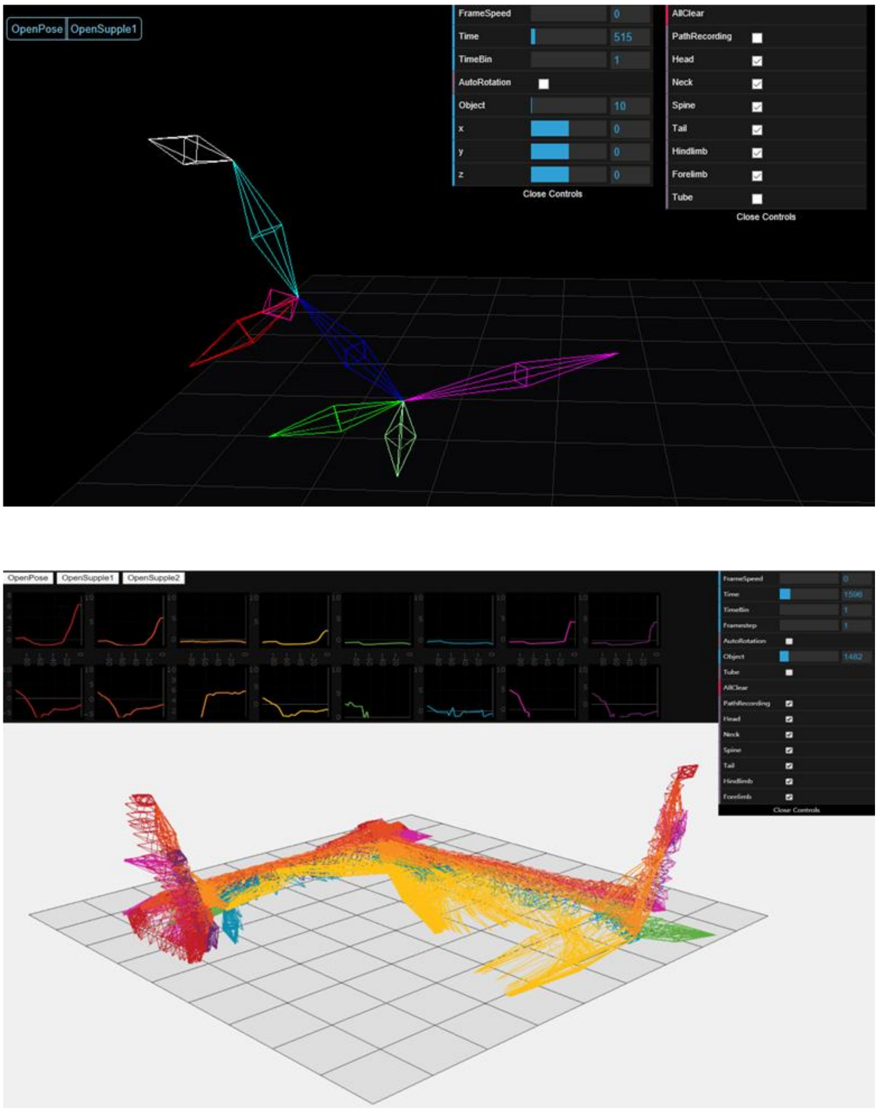


**Supplementary Figure 12: Graphical user interface (GUI) of the Web simulator in AVATAR system.**

(a) Representative image of Web simulator visualizing an action skeleton from the AVATAR coordinates file. (b) Representative image showing the stacked behavioral trajectories.

a

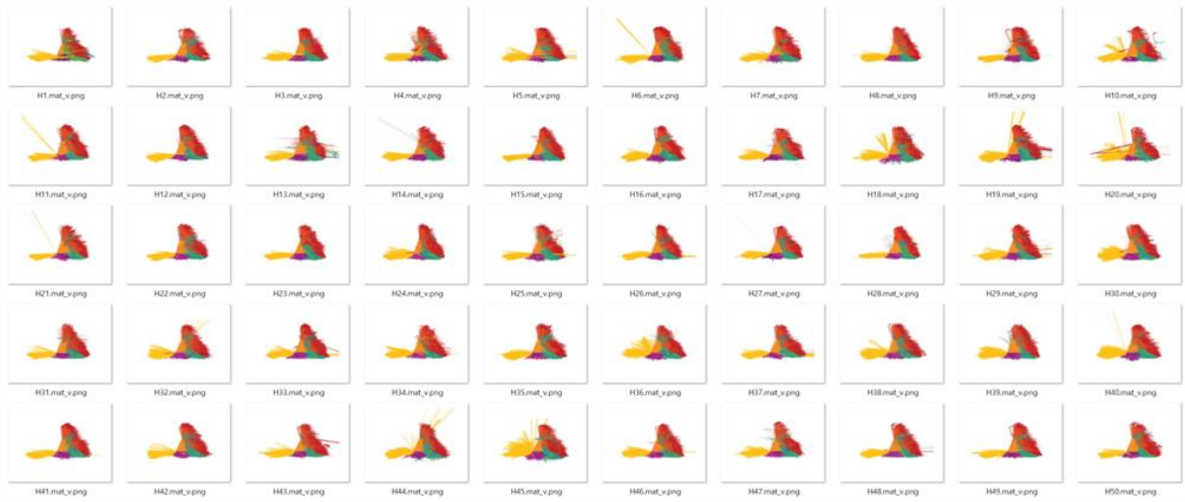

b

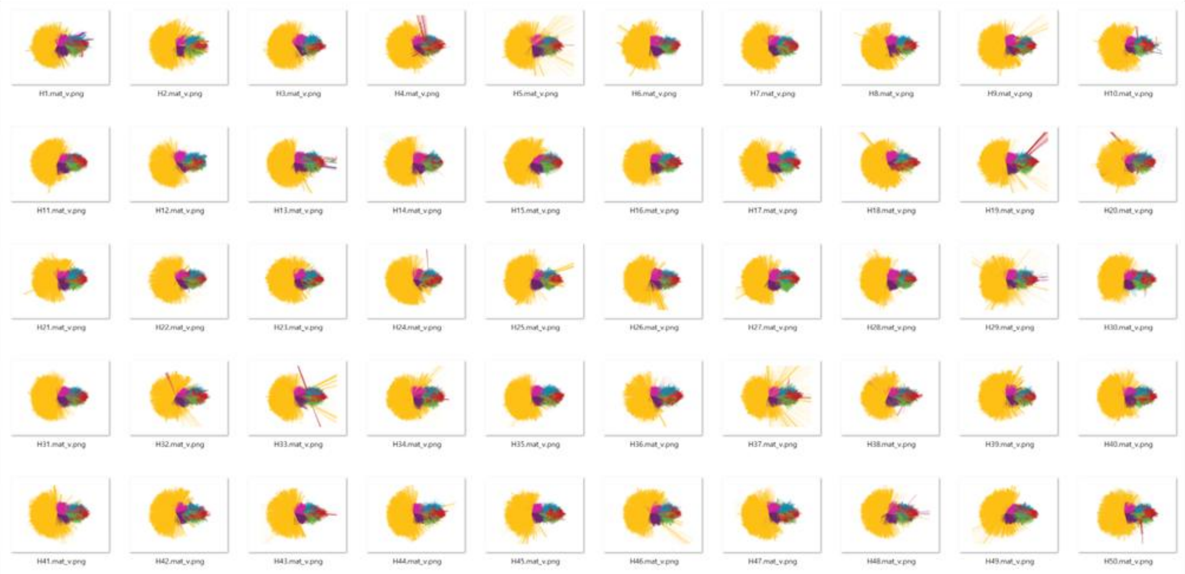

**Supplementary Figure 13: Degree of freedom of action skeleton in wild type mice.** (a-b) Stacked action skeletons representing the degrees of freedom for each joint from the lateral view (a) and top view (b). The images were obtained over 5 minutes from fifty mice, and the degree of freedom was determined by fixing the anus as the origin point.
